# The Molecular Origin of Water-Mediated Collagen Contraction

**DOI:** 10.64898/2026.03.27.713712

**Authors:** James Rowe, Peter Fratzl, Daniele Dini, Nicholas Harrison, Richard Abel, Ulrich Hansen

## Abstract

The mechanical toughness of bone and teeth relies on residual stresses generated during mineralisation, where the dehydration of collagen fibrils leads to contraction, putting the mineral phase under compression. While macroscopic stiffening of collagen upon drying is well-documented, the atomic-level structural rearrangements driving this phenomenon have remained elusive. By performing molecular dynamics simulations, we demonstrate that collagen contraction is not homogeneous but is driven by specific charged motifs. We identify a critical sequence-dependent rule for contraction: oppositely charged side chains must be separated by at least four residues to drive backbone contraction. While salt bridges can form between side chains at a distance less than four residues without perturbing the helix, those at greater distances cannot form without rupturing backbone hydrogen bonds. Consequently, dehydration forces these distant charges together, breaking local backbone structure and driving collagen contraction. These findings imply that collagen sequences are evolutionarily tuned to actively control tissue mechanics and redefines collagen as an active mechanical element rather than a passive scaffold. Furthermore, this framework provides a molecular basis for understanding mechanical failure associated with pathologies and ageing, while simultaneously opening avenues for designing bio-inspired materials with tunable pre-stress and fracture resistance.

## Introduction

Collagen is a key component of a wide variety of biological materials including, but not limited to, bone, teeth, tendon, cartilage and skin. Water is another key component in these materials, where it has been established that it directly hydrates the collagen fibrils as well as other components of the structures. It is well known that the mechanical properties of tendon and bone change upon drying, with tendon stiffening [2] and bone becoming more brittle [3], but the origin of these changes at the molecular scale is not well understood [4, 5]. Furthermore, contraction due to decreasing hydration has been implicated in the formation of residual stresses in bone and dentine [6–8]. It has been proposed that during bone mineralisation, the mineral phase replaces water, dehydrating the collagen molecules [9, 10]. This dehydration leads to contraction of the collagen, compressing the newly-formed mineral phase. These residual stresses are believed to improve the mechanical behavior of the brittle mineral phase under tensile stresses, analogous to steel beams in pre-stressed concrete, playing a key role in the impressive mechanical properties of bone and teeth [6–8].

Collagens are a diverse family of proteins, composed of at least 28 recognised types [11, 12]. Collagens are typified by a triple-helical structure, where three left-handed, polyproline II-like helices wrap around each other to form a right-handed triple helix [13]. The packing of the three strands in the triple-helix, necessitates that the side-chain of every third residue points into the center of the helix. This tight geometric constraint leads to an amino acid sequence in which every third residue is a glycine (G), the smallest amino acid [13]. Indeed, deviations from this pattern are associated with diseases such as osteogenesis imperfecta [14, 15], indicating that the resultant disruption to the structure results in catastrophic consequences for bone strength and toughness. The primary type of collagen in bones, tendons and teeth is type I [13, 16]. Collagen type-I is composed of two alpha-I chains and one alpha-II chain with the three chains forming the collagen triple helix. Collagen type-I helices pack together in a semi-regular fashion to form fibrils which have a distinctive packing along the length of the collagen helices, known as the D-period [17]. The D-period is approximately 67 nm, making it roughly 4.5 times shorter than the collagen molecule, which is approximately 300 nm. Because the D-period is not an integer divisor of the collagen length, the fibril is structured into alternating zones of high and low density, known as the overlap and gap regions, respectively [13, 17].

Due to the significant disorder within the collagen fibril, obtaining high-resolution X-ray crystallography structures of the molecular structure is difficult and does not provide sufficient detail about the interaction of the protein side-chains [18, 19]. As a result, collagen mimetic peptides (CMPs), small triple-helical molecules with sequences similar to natural collagen, have a long history as more easily characterised models of collagen molecular structure [20–22]. CMPs generally have repetitive amino acid patterns of the form glycine-proline-proline (GPP) or glycine-proline-hydroxyproline (GPO). These triplets are chosen as they lead to stable triple-helical folds and represent a significant proportion of the triplets found in naturally occurring collagen [13]. In this work, we aim to understand the behavior of different segments of the collagen type-I collagen sequence by inserting them into the center of GPP helices, forming a “guest-host” sequence. This ensures that the guest sequence is simulated in a triple-helical environment similar to that of naturally occurring collagen.

Recent experimental work has shown the sensitivity of collagen mechanical properties to changes in hydration [2, 23]. By measuring the change in D-period upon changes in humidity, X-ray scattering experiments have shown that type I collagen fibrils contract upon drying. This contraction is associated with a change in the ratio of the gap and overlap regions, suggesting that the contraction is not homogeneous across the sequence. The sequence dependence of the contraction has been explored further via experimental work utilizing collagen mimetic peptides (CMPs) [23]. These experiments showed that collagen sequences display a diverse spectrum of responses upon increasing osmotic pressure, ranging from contraction to expansion. Importantly, sequences containing charged residues were implicated as being important for contraction. However, a detailed mechanistic understanding is currently not available due to high experimental costs and the difficulty in obtaining high-resolution data at the atomic scale [24]. For example, although charged sequences are recognized as important, it is currently not known whether this applies to all charged sequences or only a subset [2, 23].

This work aims to expand on previous studies and establish the key motifs in collagen sequences that drive contraction due to dehydration. Full atom resolution molecular dynamics simulations of CMPs are performed, providing high-fidelity insights beyond what can be achieved via experiment alone. These CMPs are simulated in a high-throughput manner, providing the osmotic pressure response of a wide range of collagen sequences. By separating these CMPs based on their responses, the key features and mechanisms controlling contraction can be discovered.

We find that the behavior of collagen to decreasing hydration is dominated by specific charged regions. Counterintuitively, regions composed of sequences in which oppositely charged residues are close to each other and separated by less than 4 amino acid registers are not found to contract significantly, whereas when the distance is 4 or greater, contraction is observed. This discovery provides concrete evidence for the molecular mechanism of osmotically induced collagen contraction, and shows that the mechanical properties of collagen-based materials are directly controlled by the underlying amino acid sequence of collagen.

## Results

### Specific Charged Sequences Show Significant Contraction as they Dehydrate

Figure 2a illustrates the dependence of collagen length on hydration level for specific guest sequences. Two distinct behaviors are evident: sequences either exhibit minimal structural change or undergo significant contraction upon dehydration. Notably, sequences AKGEPGDAGAKG and NOGADGQOGAKG exhibit the most pronounced contraction. Both sequences possess oppositely charged residue pairs separated by a sequence distance of at least four residues. However, PAGPOGEAGKOG and SOGSOGPDGKTG show negligible contraction despite also containing charged pairs. This implies that the presence of charges alone is insufficient to drive contraction; rather, a critical separation distance between the pair is required. This trend holds true for the full set of tested guest sequences, as shown in Figure 2b. To capture the underlying behavior, the mean rise per residue was modeled using a quadratic fit centered at 1.05 waters/residue (*a*(*x* − 1.05)^2^ + *b*(*x* − 1.05) + *c*). Because 1.05 waters/residue represents the center of the simulated hydration range, the resulting parameters (a and b) directly describe the shape and slope of the response curve at its central point. This analysis confirms the separation of the behaviours into two general groups. Sequences located to the left of the *a* = 0 line exhibit minimal length change, generally showing only a slight expansion (≈1%) upon dehydration. This group includes the sequences LQGPOGPOGSOG, PMGPOGLAGPOG, PAGPOGEAGKOG and SOGSOGPDGKTG, which display little sensitivity to changes in hydration, as shown in Fig. 2a. Conversely, the bottom-right quadrant contains sequences exhibiting the strongest contraction, including AKGEPGDAGAKG and NOGADGQOGAKG. These sequences display a positive gradient, indicating elongation upon hydration, coupled with negative curvature, suggesting saturation toward a constant length at high water content. This plot confirms a clear division in behaviour based on the distance between oppositely charged pairs. The stable group generally comprises neutral sequences and those with charge separations of ≤3 residues, while the contracting group consists of sequences with charge separations *>* 3 residues. A small subset of sequences with charge separations *>* 3 deviate from this trend, appearing in the stable group with no significant contraction. To explain this discrepancy, a representative example, ERGFOGERGVQG (whose curve is also shown in Fig. 2a), is analyzed in detail in the subsequent section.

**Figure 1:**
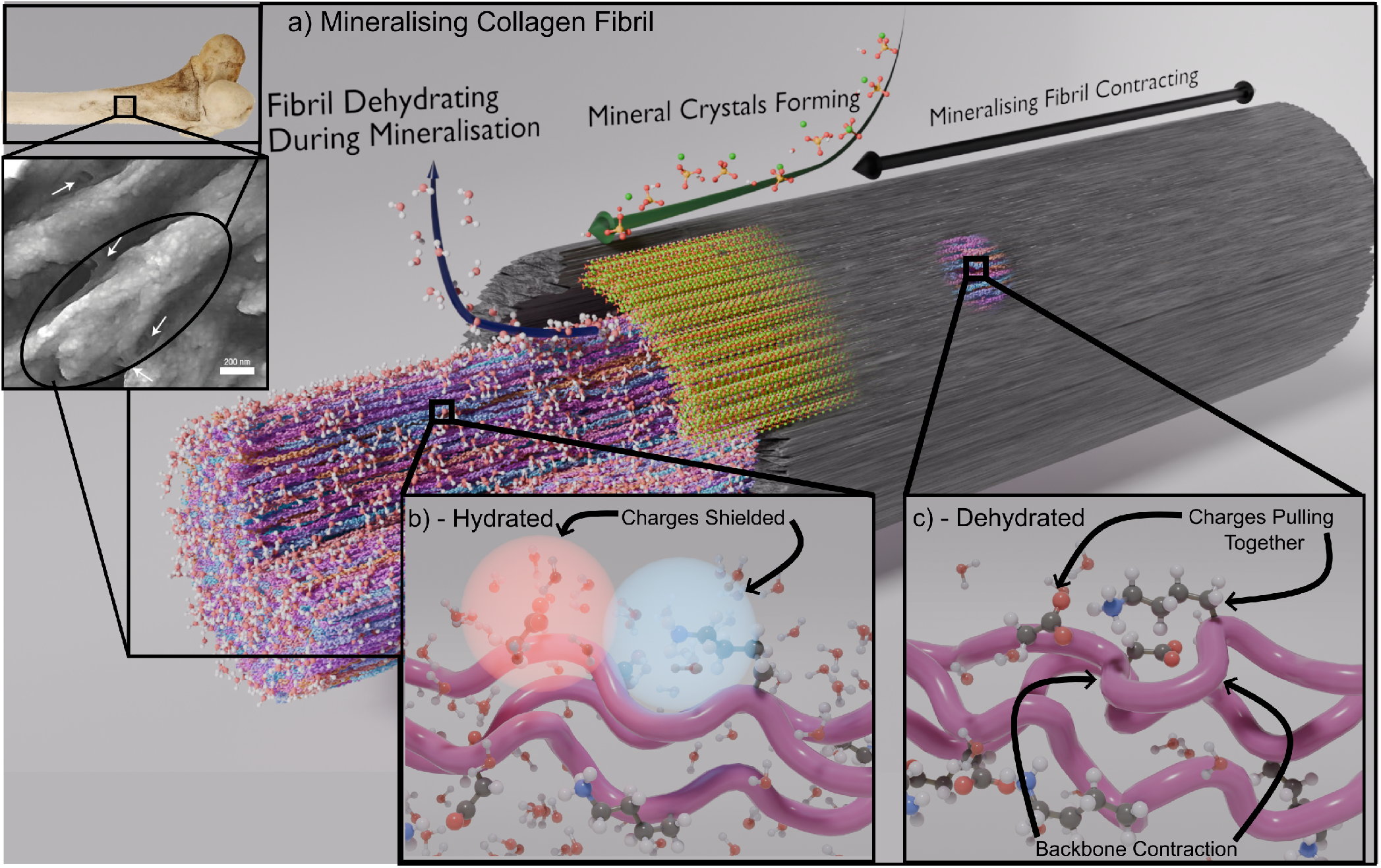
Demonstration of the mechanism of osmotic-pressure mediated residual stress formation in bone. Top left: An image of a femur and an AFM image of mineralised collagen I fibrils. AFM image reprinted from [1]. (a): A semi-mineralised collagen fibril. As the fibril is mineralised, water is displaced by mineral crystals dehydrating the fibril. (b): A simulation snapshot of a collagen-mimetic peptide in high-hydration conditions. Water molecules solvate the charged side chains shielding them from each other. (c): A simulation snapshot of a collagen-mimetic peptide in low-hydration conditions. The lack of water causes the attraction between the charges to increase and they pull towards each other, causing the backbone to contract.

**Figure 2:**
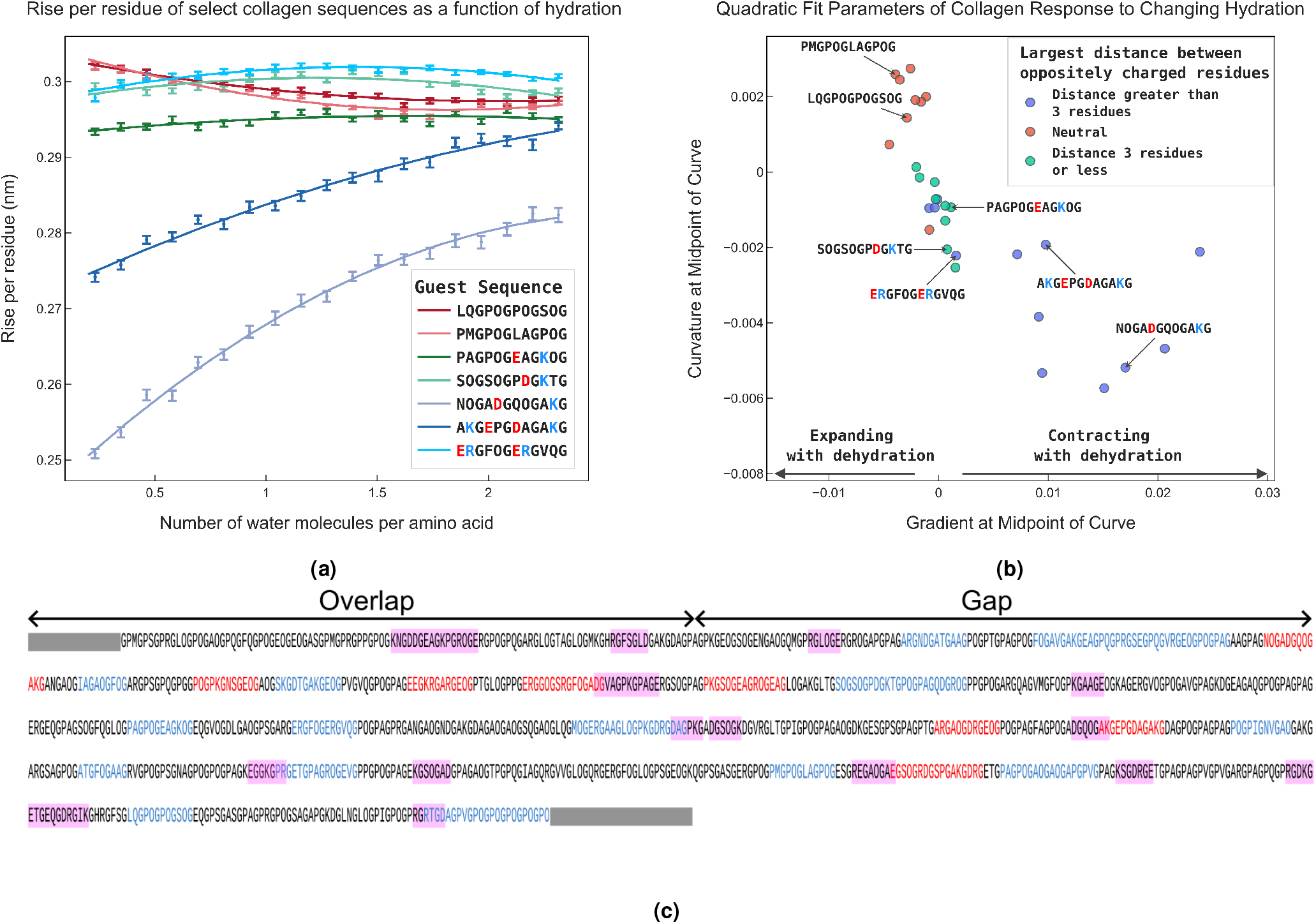
(a): Rise per residue of selected collagen guest sequences due to changes in hydration. The points represent the mean values of the rise per residue from MD simulations, whilst the error bars are the standard error in the rise per residue from the simulations. The solid lines represent quadratic fits to the data. In the legend, negative residues are in red font and positive residues are shown in blue font. Rise per residue curves can be found for all other tested sequences in S1. (b): Coefficients from fitting a quadratic of the form *a*(*x* − 1.05)^2^ + *b*(*x* − 1.05) + *c* to the mean rise per residue at each hydration level for each sequence. The quadratic is fitted around 1.05 water molecules per amino acid as this is the center of the range of water molecules per amino acid tested. This plot shows *a* (the curvature) against *b* (the gradient) for each guest sequence simulated. Sequences with greater than 3 residues between oppositely charged residues (blue) generally contract upon dehydration (positive gradient). Conversely, neutral or sequences with a distance of 3 residues or less between oppositely charged residues (red/green) slightly expand upon dehydration (negative gradient). (c): The location of the tested sequences in the COL1A1 *Homo sapien* sequence [25]. Tested sequences that were found to contract upon dehydration (gradient *>* 0.005 nm) are colored in red, whereas sequences that expanded or showed little change are colored in blue. Other potentially contracting sequences are highlighted in pink. These sequences were found by looking for regions with a distance separating oppositely charged residues of between 4 and 7 residues. The non-helical telopeptide regions are represented by gray rectangles.

The location of all the tested sequences within the collagen fibril is shown in Fig. 2c. In this figure, tested sequences that were found to contract upon dehydration (gradient *>* 0.005 nm) are colored in red, whereas sequences that expanded or showed little change are colored in blue. Other potentially contracting sequences are highlighted in pink. These sequences were chosen based upon whether they contained oppositely charged residues separated by a distance between 4 and 7 residues. Following on from the observed patterns in Fig. 2b, we believe that this a necessary but not sufficient condition for the region to be contracting. By examining the distribution of the tested (red) and predicted (pink) sequences, regional behaviors emerge across the fibril. For example, many of the contracting sequences in the overlap region appear to be confined to a single band, whereas in the gap region contracting sequences appear to be much more uniformly distributed. These differences in response between the two regions is supported by experimental observation which found greater contraction in the gap region [2]. Key sequences are analysed further in the next sections.

### Salt Bridge Network Analysis

To investigate the molecular origin of contraction further, the PAGPOGEAGKOG, AKGEPGDAGAKG and ERGFOGERGVQG sequences were analysed in greater detail. These sequences were chosen as they all contain some number of oppositely charged side chains, but they show different responses. Salt bridges are common interactions in proteins that occur between oppositely charged amino acid side chains, arising from a combination of electrostatic and hydrogen-bonding interactions. For these three sequences, the probability of each salt bridge was calculated at different hydration levels. Using these probabilities, salt bridge networks were constructed and are shown in Fig. 3 for two hydration levels. In these networks, positively charged side chains are represented as blue nodes and negatively charged side chains as red nodes. A connection is drawn between nodes if a salt bridge occurs more than 5% of the time. The thickness and color of each connection reflect the frequency of salt bridge formation, with thicker connections indicating higher probabilities. To aid visualization, only one layer of three helices is shown.

**Figure 3:**
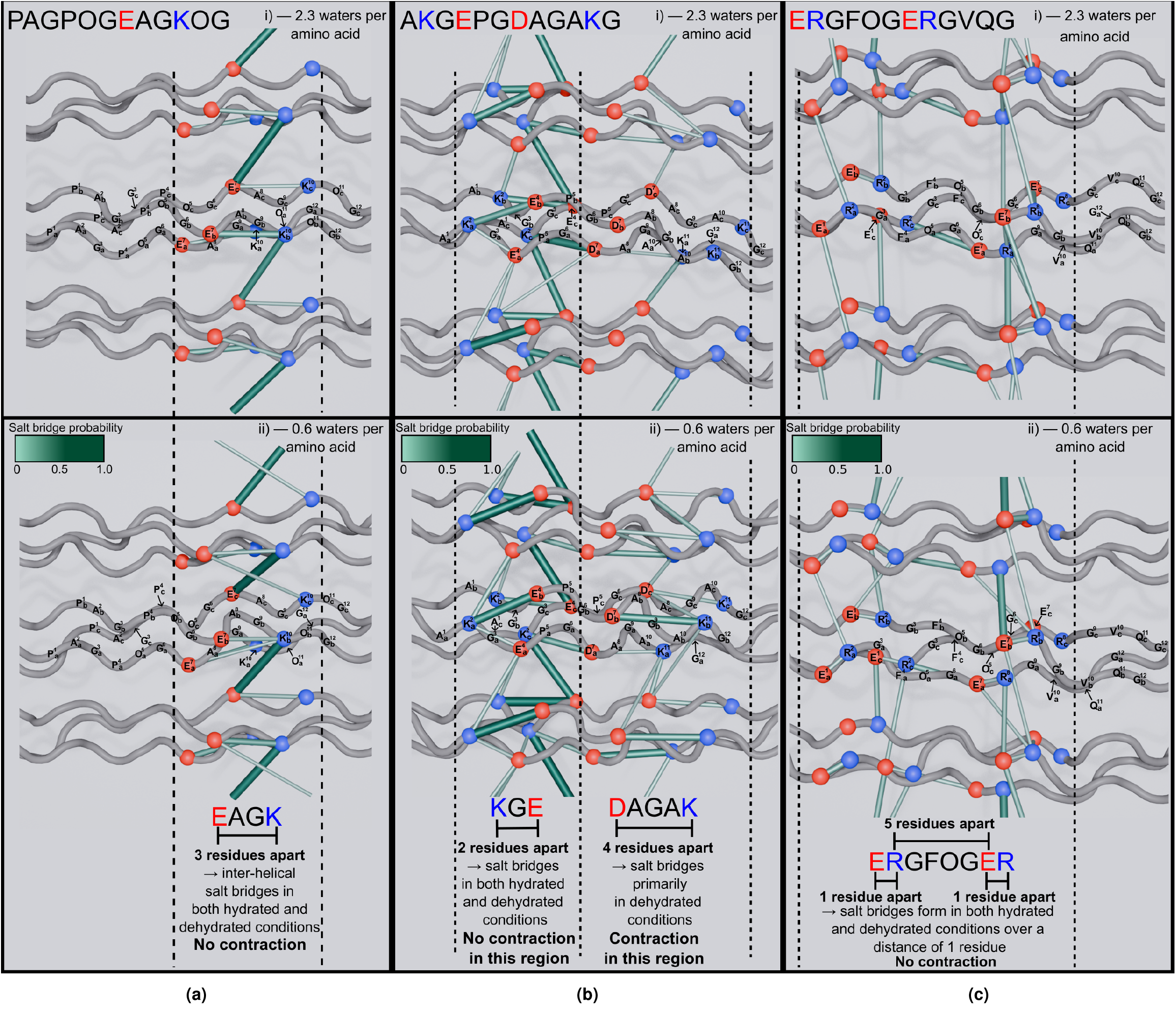
Salt Bridge networks of three guest sequences in two different hydration conditions. (a): PAGPOGEAGKOG. (b): AKGEPGDAGAKG. (c): ERGFOGERGVQG. The blue spheres mark the location of the C_*α*_ atoms of the positively charged residues, with the red spheres marking the location of the C_*α*_ marking the location of the negatively charged residuces. The location of the guest-sequence residues are marked on the middle helix, with the subscript denoting the chain and the superscript denoting the location within the guest-sequence. A green rod is shown between the spheres if a salt bridge was observed between those residues more than 5% of the time. Thicker and darker rods correspond to salt bridges of higher probability.

#### Non-Contracting Charged Sequence: PAGPOGEAGKOG

The salt bridge networks for the PAGPOGEAGKOG sequence with high hydration (2.3 water molecules per amino acid) and low hydration (0.6 water molecules per amino acid) are shown in Fig. 3a. Across both hydration levels, a significant number of salt bridges appear. These include both intra-helical (within a helix) and inter-helical (between adjacent helices) salt bridges. The intra-helical salt bridges are mostly between groups on the same chain and therefore span a distance of three amino acid registers (as there are three registers between the E and K in the EAGK region on a chain). This sequence does not show significant contraction, as seen from its curve in Fig. 2a. The fact that intra-helical salt bridges can form over a distance of three registers in both high and low hydration without contracting the helix, implies that salt bridges must form over longer distances for contraction to be seen.

#### Contracting Charged Sequence: AKGEPGDAGAKG

The AKGEPGDAGAKG sequence was selected for further analysis as it contains both an oppositely charged pair at a distance of 2 (between the K and E in KGE), and an oppositely charged pair at a distance of 4 (between the D and K in DAGAK). This sequence gives insights into the behaviour of oppositely charged pairs at different distances. Across both hydration levels shown, prominent salt bridges appear in the AKGEPG region of the molecule. As with PAGPOGEAGKOG, these include both intra-helical and inter-helical interactions. Because these salt bridges already occur with high probability even under high hydration, their likelihood does not increase substantially upon dehydration. In contrast, the DAGAK region shows few salt bridges under high hydration. As water is removed, both the number and strength of salt bridges in this region increase, indicated by an increase in the number and thickness of connections. This trend suggests that the formation of salt bridges in the DAGAK region drives the overall contraction observed during dehydration. To further explore the difference in salt bridge formation between the two regions, snapshots of the charged side chains of the AKGEPGDAGAKG sequence under two different hydration conditions are shown in Fig. 4a and Fig. 4b. Salt bridges between the K and E side chains within the AKGEPG region are observed at both high and low hydration. Due to the spatial closeness of these side chains, the salt bridge can form without having to disrupt the helix backbone. On the other hand, in high hydration the D and the K within the DAGAK region are separated from each other. In low hydration, salt bridge formation between the D and K side chains is observed, as was ascertained from visualisation of the salt bridge network. Significant disruption in the backbone structure around this region is observed, implying that the backbone must be disrupted in order for the salt bridges in this region to form.

**Figure 4:**
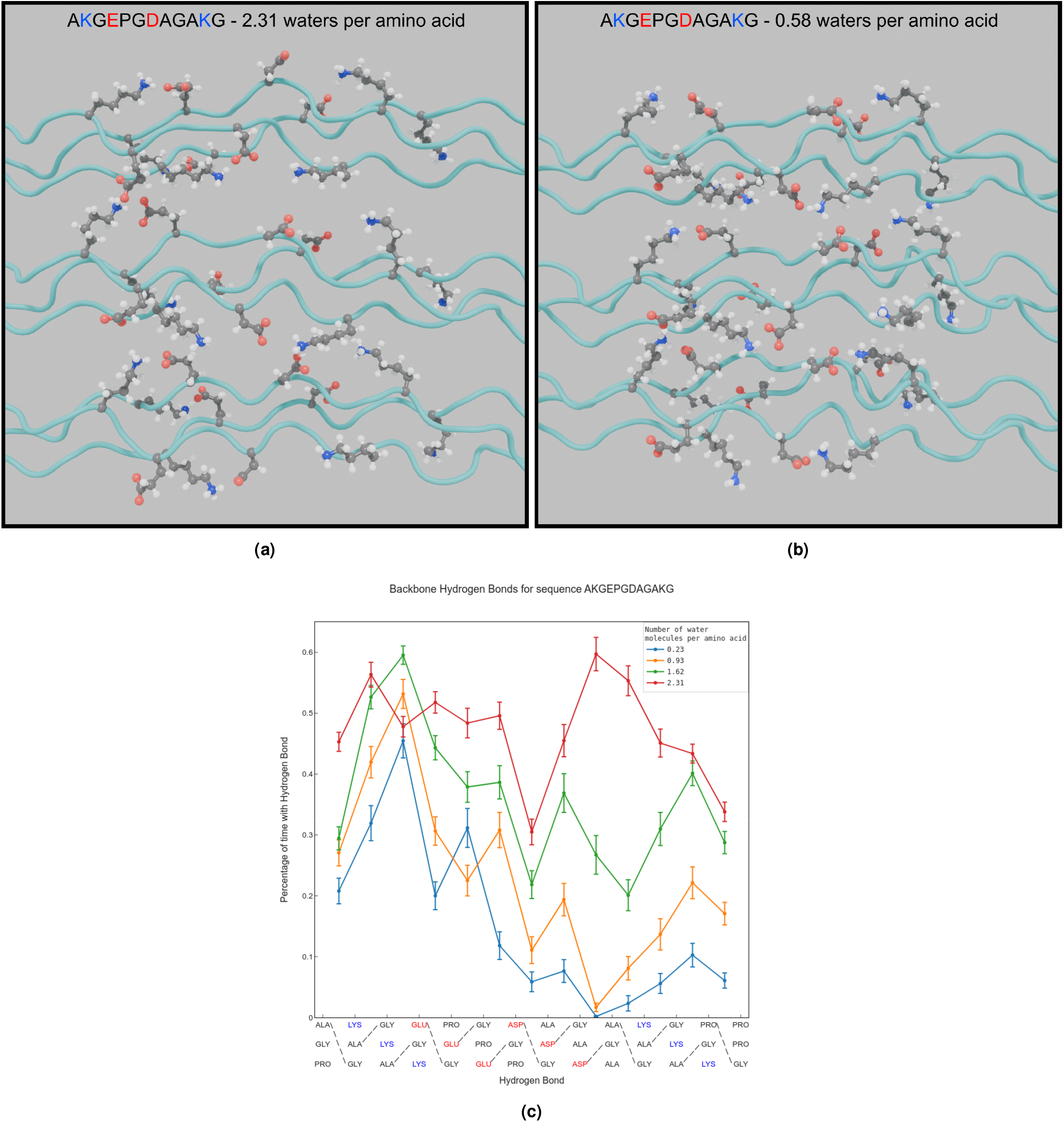
(a) and (b): Snapshots from simulations of the contracting guest sequence AKGEPGDAGAKG in two different hydration conditions. In high hydration conditions (a), the charges in the DAGAKG region are well solvated and do not form salt bridges. In low hydration conditions (b), the salt bridges form, leading to significant disruption of the backbone. (c): Backbone hydrogen bond probability for AKGEPGDAGAKG in the guest sequence region. The probabilities are shown for 4 different levels of hydration. In the region around the DAGAKG sequence there is significant disruption of the backbone hydrogen bonds.

#### Sequence not Following Distance Trend: ERGFOGERGVQG

The salt bridge network for the ERGFOGERGVQG sequence in both high and low hydration is shown in Figure 3c. This sequence was analyzed further because there is a distance of five residues between an oppositely charged E-R pair (and another pair at a distance of seven residues), yet no contraction is observed. This sequence belongs to the group of blue points that are shown to not significantly contract in Fig. 2b. The salt bridge network for this sequence indicates that, at both hydration levels, the majority of salt bridges form within the two ERG sections, with few salt bridges connecting these regions. For each helix, there is one salt bridge that connects the two regions. This salt bridge appears at all hydration conditions tested and forms between the first R on the third chain (marked 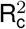) and the second E on the first chain (marked 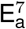). Since the three collagen chains that compose a helix are staggered by one residue relative to one another, these two side chains are only three registers apart rather than five. The fact that this salt bridge does not cause contraction lends further support to the view that a distance greater than three registers is required for a salt bridge to cause contraction.

### Backbone H-Bonding Analysis of the Contracting AKGEPGDAGAKG Sequence

To better understand the driving forces of contraction and how contraction alters the collagen structure, the hydrogen bonding of the backbone for the AKGEPGDAGAKG sequence was analyzed. This analysis is shown in Fig. 4c. This analysis reveals that there is a significant decrease in the hydrogen bonding of the backbone as the sequence is dehydrated. Importantly, this disruption is localised around the DAGAK region, with hydrogen bonding only slightly decreasing around the KGE region. At very low hydration, the hydrogen bond in the center of the DAGAK region is almost never observed. This indicates that when the oppositely charged residues are close, salt bridges can form without disrupting the backbone and causing contraction. However, when the charged residues are further apart the backbone hydrogen bonds need to broken before the salt bridges can form.

## Discussion

Our data reveals that water-mediated contraction develops in regions where oppositely charged residues are separated by at least four positions along the sequence (i.e. one residue at position *i* and the other at *i* + 4 or beyond). As collagen dehydrates, reduced solvation weakens the electrostatic shielding of these charges, increasing the effective attraction between positively and negatively charged side chains [26, 27]. When the residues are close together (fewer than four positions apart), salt bridges can easily form without distorting the backbone. However, when residues are farther apart, salt-bridge formation requires the backbone to contract, bringing the side chains into contact. This backbone compaction leads to local disruption and contributes to residual stress as the system dehydrates. It is well established that salt-bridge formation is essential for stabilizing collagen and collagen-mimetic peptides [26, 28, 29]. These stabilizing salt bridges are typically interchain and involve residues separated by three or fewer positions. This work reveals a novel mechanism: at larger separations (four or more positions), salt bridges can also form, but only under low-hydration conditions, driving backbone contraction and contributing to residual stress formation.

The critical distance of 4 residues can be rationalised using data on salt bridges from protein databases. By analyzing protein databases, Donald et al. found that positively charged side chains can occupy different geometries in salt bridges [30]. They calculated the mean C*α*–C*α* distance for the different possible salt bridge side chain geometries. For the four possible pairs of salt bridges that can form between K or R and E or D, the largest mean distances were formed from the geometries of K-trans – D, K-trans – E, R-end-on – D, and Rend-on – E. These salt bridges were found to have mean C*α*–C*α* distances of 9.5 Å, 9.6 Å, 10.2 Å, and 10.9 Å, respectively [30]. From these pairs, we can obtain a lower and upper bound for the maximum backbone distance over which a salt bridge can form without requiring backbone contraction. Using a rise-per-residue for the collagen backbone of 3.0 Å (as was found to be the rise-perresidue for hydrated collagens; see Fig. 2a), the lower and upper bounds are estimated to be 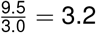 residues and 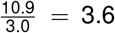 residues respectively. This means any salt bridge formed between a pair of K or R and E or D can span at most 3 residues between the C_*α*_ atoms without requiring contraction. If the separation exceeds 3 residues, the rise-per-residue must decrease to allow the salt bridge to form. Histidine (H) is another amino acid that can be positively charged at physiological pH. Histidine is excluded from our analysis as its residues are rare in collagens compared to lysine (K) and arginine (R) [31].

A handful of sequences were found not to follow the simple trend observed for most of the sequences. These sequences correspond to the blue points located in the top-left of Fig. 2b and do not show contraction when they were expected to, based on the simple analysis of the largest distance between oppositely charged amino acid side chains. The salt bridge network for one of these sequences, ERGFOGERGVQG, was previously shown in Fig. 3c. The network demonstrates that, when possible, side chains tend to form salt bridges with other side chains in closer spatial proximity. Because adjacent, oppositely charged side chains prevent salt bridges from forming between distant residues, the expected contraction is not observed. A similar explanation applies to all the other outlier sequences: in each case the arrangement of the charged side chains is such that each charge can form salt bridges without having to form salt bridges that lead to contraction.

In all simulations, CMPs were packed adjacent to others with the same sequence, creating a potentially unrealistic environment compared to naturally occurring collagen fibrils. In nature, the staggered arrangement of the collagen fibril results in diverse adjacent sequences, a distinction that may be particularly significant for charged motifs where salt bridges form both within and between helices (Fig. 3). Within a fibril, the availability of different neighboring helices could alter the behavior of these sequences. This may be relevant to the behavior of the previously mentioned outlier sequences, which resist contraction by forming short-range salt bridges. As seen in the ERGFOGERGVQG network (Fig. 3c), many of these connections are inter-helical. If these helices were instead surrounded by neutral side chains, long-range salt bridges might form more readily, potentially leading to contraction. Consequently, further investigation into the specific nature of charged connections within collagen fibrils is required.

Analysis of the backbone hydrogen bonding structure shows that backbone hydrogen bonding is disrupted around the contracting region. There is clear competition between salt bridge formation and the backbone hydrogen bonds. At high hydration the charged side chains are well solvated, stabilising the charges. It can be inferred that any gain in free energy from forming the salt bridge is counteracted by the free energy loss caused by the breaking of the backbone hydrogen bonds. As the structure becomes dehydrated, the packing of the helices becomes tighter, reducing the solvent in the interhelical space. The loss of solvent in the inter-helical space decreases the solvation of the charged side chain and increases the free energy gain from forming the salt bridge. As a result, contraction starts to occur as the backbone hydrogen bonds start to be broken.

In this work, guest sequences were chosen from the *Homo-Sapien* COL1A1 and COL1A2 sequences to test the response of a realistic set of collagen sequences. Although the sequences were taken from a specific collagen type, this work was focused on finding the underlying motifs that drive contraction caused by dehydration and is applicable to any protein showing the same triple helical structure. Consequently we believe our results contribute to the understanding of a number of collagen types. Indeed osmotic pressure changes are known to play a crucial role in the mechanical behavior of cartilage, which is primarily composed of collagen with the most common type being type II [32–34]. Equipped with the understanding developed in this paper, further analysis of the charge distribution and salt bridge networks in type II collagen could lead to new insights into stress formation in cartilage.

In conclusion, this work establishes a direct, mechanistic link between the primary amino acid sequence of collagen and the macroscopic material properties of structural tissues. By uncovering the critical role of charge spacing in controlling backbone contraction, we move beyond viewing collagen as a passive scaffold and show that the material properties of a patient’s bone, cartilage, and skin are intrinsically defined and actively controlled by their underlying collagen genetic sequence. Specific salt bridges serve as molecular “switches”, generating residual stresses, a fundamental requirement for bone toughness. Consequently, this framework not only provides a molecular basis for understanding mechanical failure associated with pathologies and ageing [35], but also offers possibilities for designing bio-inspired materials with tunable pre-stress and superior fracture resistance.

## Methods

### Constructing Collagen-Mimetic Peptides

Collagen-Mimetic Peptides (CMPs) were constructed using the Triple HElical BUilder SCRipt (THeBuScr) [36, 37]. Each CMP chain was constructed by inserting a guest sequence between six PPG triplets. Each chain was capped by neutral acetyl (Ace) and amide (Nme) capping groups at the N-terminal and the C-terminal respectively. A variety of guest sequences were simulated ranging in size from 12 to 18 amino acids. The majority of CMPs were homotrimers, however some heterotrimers were also tested. For heterotrimers, an AAB arrangement of the chains was chosen. Although the arrangement of the chains in type-I collagen is not currently known, this arrangement was chosen based upon recent work [38]. The guest sequences were taken from the triple-helical region of the *Homo sapien* collagen type-I alpha-I and alpha-II chains.

### Simulation Details

To build the simulation box, each CMP was replicated 9 times, and the 9 helices were packed together in a hexagonal pattern, inspired by the hexagonal packing exhibited in the overlap region of collagen type-1 fibrils. Images of this simulation setup can be found in Fig. S2. Varying numbers of TIP3P water molecules [39] ranging from 0.23 to 2.31 water molecules per amino acid residue were added to these boxes using the GROMACS solvate program [40]. This range of water molecules was chosen as they give lateral distances between helices similar to the distance between helices in collagen fibrils. This range of water molecules is also similar to estimates for the number of waters per amino acid from X-ray scattering experiments [23]. For each number of water molecules, 10 replicas were simulated to improve statistical sampling. To ensure each replica began with a distinct molecular arrangement, water molecules were added using independent random distributions. The collagen molecules were modelled using the Amber99SB*-ildnp force field, which is well established for simulating collagen [41–44]. Using the GROMACS program [40], each replica was subject to the following procedure: energy minimisation, 1 ns NVT equilibration with the protein heavy atoms restrained, 1 ns NPT equilibration with the protein heavy atoms restrained, 2 ns NPT equilibration without the protein heavy atoms restrained and finally a 5 ns production run in NPT conditions. In total, across the 10 replicas, 50 ns of production simulation were performed for each hydration condition per CMP. In the constant temperature simulations, a stochastic velocity-rescaling thermostat set to 310 K was used with a time constant of 0.1 ps [45]. A 1 bar semi-isotropic stochastic barostat with a time constant of 5 ps [46] was used in the NPT simulations, with the x-y directions scaled together in order to conserve the hexagonal packing of the molecules (see Fig. S2 for images of the simulation box). The z direction was allowed to scale independently. All bonds involving hydrogen atoms were constrained to their initial length [47, 48]. This permitted the use of a 2 fs timestep. Long-range electrostatic interactions were calculated via the Particle Mesh Ewald (PME) method with a 1.0 nm real-space cutoff [49, 50]. Van der Waals interactions were truncated at 1.0 nm with dispersion corrections applied for energy and pressure.

### Analysis

The simulations were analysed using MDAnalysis [51, 52].

#### Rise-per-residue Analysis

For a given guest sequence and hydration level, the average length along the helical axis of the guest sequence region was measured for each replica. This length was divided by the number of residues in the guest region to obtain the rise per residue. This rise per residue was then averaged over each replica to obtain a plot of rise per residue against hydration for each guest sequence and obtain standard error estimates (Fig. 2a). A quadratic equation of the form *a*(*x* − 1.05)^2^ + *b*(*x* − 1.05) + *c* was fitted to the rise per residue vs hydration data for each sequence. The parameters *a* and *b* were used to obtain the curvature and gradient of the curve at waters/amino acid.

#### Salt Bridge Networks

A positively charged side chain (K or R) and a negatively charged side chain (E or D) were defined as having formed a salt bridge if they followed the condition stated by Barlow and Thornton [53]. This condition requires that there is a maximum distance of 4 Å between a nitrogen atom on the positive side chain and an oxygen atom on the negative side chain. The probability of a given salt bridge was calculated by the fraction of the frames across all replicas that the salt bridge was present. These probabilities were used to construct a network between positive and negative amino acid residues using the NetworkX [54] Python module. A node was created for each amino acid residue and edges were created between the nodes if the probability of a salt bridge between those residues was greater than 5%. Each edge was assigned a weight equal to the probability of observing a salt bridge between the two residues. These networks were visualised using Blender with the MolecularNodes [55] plugin, with edges of greater weight given thicker lines and darker colors.

## Supporting information

This file contains all supplemental figures and tables

