## Supplementary material for "The Molecular Origin of Water-Mediated Collagen Contraction": This file contains all supplemental figures and tables

February 2026

#### 1 Tables of Quadratic Fit Parameters For All Tested Sequences

##### 1.1 Homotrimers

| Guest Sequence | $a$ (nm) | $b$ (nm) | $c$ (nm) | Raw Data & Fit |
|---|---|---|---|---|
| IAGAOGFOGLQG | 0.304 | -0.00450 | 0.00073 | Fig. S1a |
| PMGPOGLAGPOG | 0.298 | -0.00394 | 0.00259 | Fig. 2a (Main text) |
| POGPIGNVGAOG | 0.297 | -0.00354 | 0.00245 | Fig. S1a |
| LQGPOGPOGSOG | 0.299 | -0.00289 | 0.00144 | Fig. 2a (Main text) |
| GPOGPOGPOGPO | 0.288 | -0.00257 | 0.00274 | Fig. S1a |
| POGPOGPOGPOG | 0.290 | -0.00213 | 0.00191 | |
| RTGDAGPVGPOG | 0.299 | -0.00204 | 0.00013 | |
| VRGEOGPOGPAG | 0.301 | -0.00174 | -0.00014 | |
| PAGPOGAOGAOGAOGPVG | 0.296 | -0.00159 | 0.00187 | Fig. S1b |
| PPGPPGPPGPPG | 0.290 | -0.00114 | 0.00200 | |
| ATGFOGAAGVVG | 0.302 | -0.00084 | -0.00153 | |
| PQGPRGSEGPQG | 0.298 | -0.00027 | -0.00072 | |
| PRGETGPAGROGEVG | 0.298 | -0.00010 | -0.00072 | |
| FOGAVGAKGEAG | 0.303 | 0.00061 | -0.00129 | |
| SOGSOGPDGKTG | 0.300 | 0.00077 | -0.00205 | Fig. 2a (Main text) |
| PAGPOGEAGKOG | 0.295 | 0.00115 | -0.00093 | |
| ARGNDGATGAAG | 0.300 | 0.00153 | -0.00253 | Fig. S1c |
| ERGFOGERGVQG | 0.302 | 0.00160 | -0.00221 | Fig. 2a (Main text) |
| PKGSOGEAGROGEAG | 0.290 | 0.00716 | -0.00218 | Fig. S1c |
| AKGEPGDAGAKG | 0.284 | 0.00976 | -0.00192 | Fig. 2a (Main text) |
| EEGKRGARGEOG | 0.273 | 0.01510 | -0.00573 | Fig. S1c |
| NOGADGQOGAKG | 0.269 | 0.01703 | -0.00519 | Fig. 2a (Main text) |
| ERGGOGSRGFOGADG | 0.262 | 0.02062 | -0.00468 | Fig. S1c |
| AEGSOGRDGSOGAKGDRG | 0.255 | 0.02382 | -0.00212 | |

Table S1: Table of fit parameters for all the tested homotrimers from fitting a quadratic curve of the form $a(x-1.05)^2 + b(x-1.05) + c$ to the plots of rise-per-residue against number of water molecule per amino acid residue. The sequences are sorted by increasing $b$ value. Positive residues (K and R) are colored in blue whilst negative resiudes (E and D) are colored in red. The raw data and curves can be found in the figure referenced in the last column.

#### 1.2 Heterotrimers

| Chain 1 & 2 Guest Sequence | Chain 3 Guest Sequence | $a$ (nm) | $b$ (nm) | $c$ (nm) | Raw Data & Fit |
|---|---|---|---|---|---|
| SKGDTGAKGEOG | SKGESGNKGEOG | 0.303 | -0.00086 | -0.00096 | Fig. S1d |
| POGPAGQDGROG | PVGLPGIDGRPG | 0.297 | -0.00036 | -0.00027 | |
| LOGPKGDRGDAG | IOGGKGEKGEOG | 0.303 | -0.00034 | -0.00094 | |
| MOGERGAAGLOG | LOGERGAAGIOG | 0.302 | 0.00061 | -0.00089 | |
| ARGAOGDRGEOG | PRGSOGERGEVG | 0.285 | 0.00913 | -0.00384 | |
| POGPKGNSGEOG | ATGARGLVGEOG | 0.288 | 0.00944 | -0.00533 | |

Table S2: Table of fit parameters for all the tested heterotrimers from fitting a quadratic curve of the form $a(x-1.05)^2 + b(x-1.05) + c$ to the plots of rise-per-residue against number of water molecule per amino acid residue. The sequences are sorted by increasing $b$ value. Positive residues (K and R) are coloured in blue whilst negative residues (E and D) are coloured in red. An AAB ordering of the chains was chosen meaning the first two chains had the same sequence and the third chain had a different sequence. The chain 1 and 2 sequence was taken from the *Homo sapien* COL1A1 sequence with the chain 3 sequence taken from the corresponding section of the COL1A2 sequence. The raw data and curves can be found in the figure referenced in the last column.

### 2 Rise-per-Residue Plots

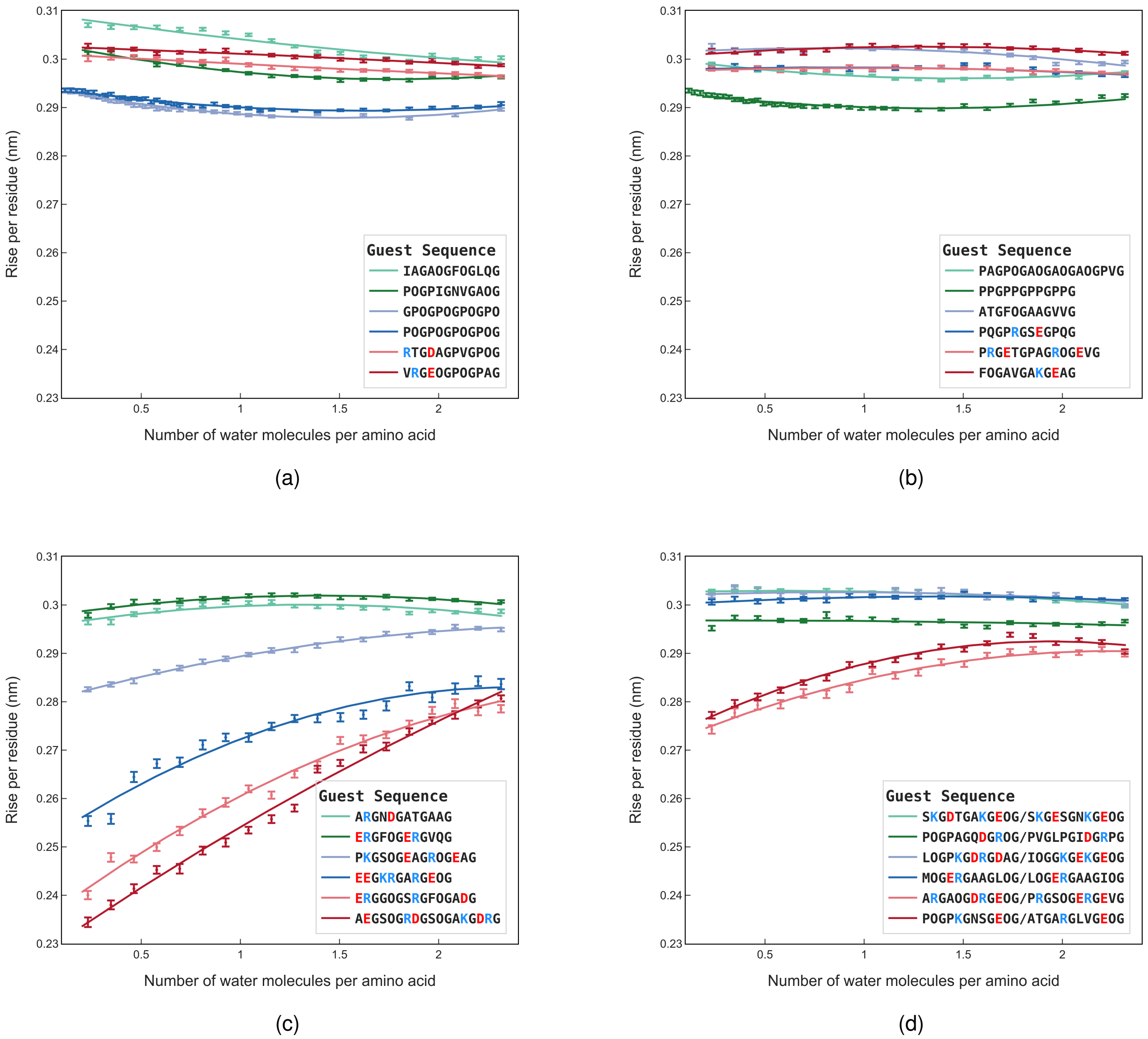


Figure S1: Plots of rise-per-residue against number of water molecules per amino acid. The marks show the raw data with the lines showing quadratic fits to this data. In Fig. (d), heterotrimers are plotted (see Table S2). In the legend, the forward slash (/) delineates the two sequences.

#### 3 Simulation Setup

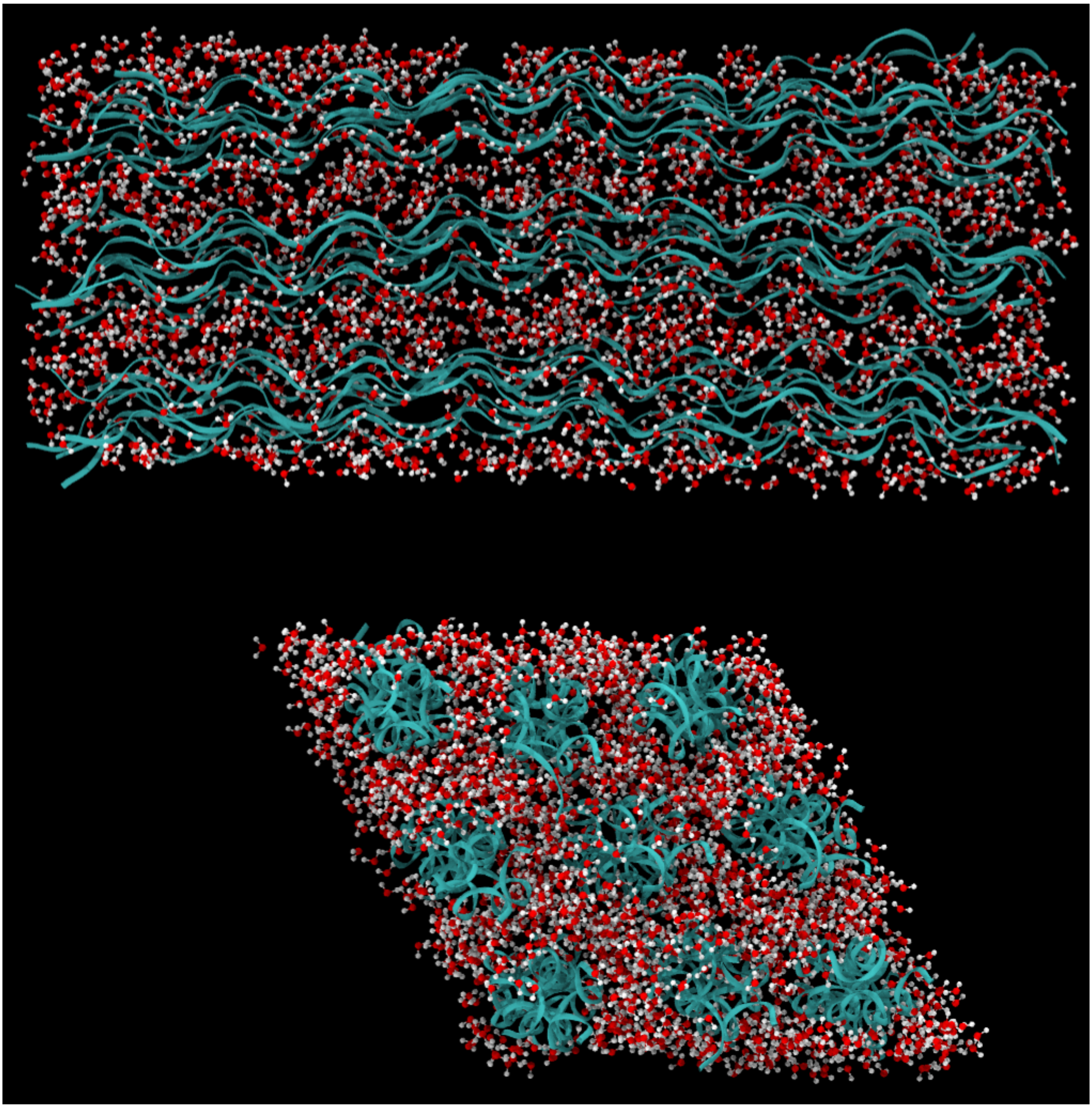

Figure S2: Top: side-on and Bottom: front-on snapshots of the simulation box. 9 identical helices were stacked in a hexagonal manner and simulated under periodic boundary conditions.
